# Effect of Electroacupuncture Pretreatment on Cognitive Impairment and Mitophagy in Aging Model Rats and its mechanism

**DOI:** 10.1101/2024.03.09.584137

**Authors:** Biyong Liu, Tiantian Tan, Jianmin Liu, Zhijie Li, Qunhu Feng, Su Qiu, Chengkai Xiong, Qing Liu, Jialin Li, Yihong Li

## Abstract

**Objective:** Investigate the effects of "Shuanggu Yitong" EA pretreatment on cognitive impairment, mitochondrial function, and mitophagy in aging model rats, and to analyze the related mechanisms.

**Methods:** Forty 3-month-old male SD rats were randomly divided into a blank group, a model group, an EA group, and a sham EA group, with 10 rats in each group. And the Morris water maze test was performed after the intervention. HE staining, to observe the morphological changes of the hippocampus of the model rats. Nissl staining was used to observe the changes in the number of hippocampal neurons in rats, Western Blotting (WB) was used to observe the expression of endogenous PTEN-induced hypothetical kinase 1 (PINK1) and human Parkinson’s protein 2 (Parkin) in the hippocampus, spectrophotometry was employed to detect the activity of respiratory chain complex I in the hippocampus of the model rats, and flow cytometry was utilized to detect hippocampal mitochondrial membrane potential (MMP) and hippocampal mitochondrial permeability transition pore (MPTP) opening.

**Results:** "Shuanggu Yitong" EA pretreatment relieved the cognitive impairment induced by D-galactose in subacute aging model rats. The mechanism of "Shuanggu Yitong" EA pretreatment in the improvement of cognitive impairment of subacute aging model rats may be related to the enhancement of Pink1/Parkin mediated mitophagy and the timely removal of accumulated abnormal mitochondria, thus improving mitochondrial function.

**Conclusion:** "Shuanggu Yitong" EA pretreatment can significantly improve the cognitive impairment induced by D-galactose in subacute aging model rats.

## Introduction

In recent years, the number of elderly people has continued to increase, and the proportion of the global elderly population will reach 22% by 2050[1]. The process of aging is accompanied by decline of various functions, including learning, memory, cognition, etc. The elderly are potential cognitive impairment patients, and likely to develop Alzheimer’s disease(AD). That the number of people with AD will continue to increase in the next decade, which will negatively affect society and individuals[2]. With the acceleration of global population aging, the demand for the prevention and treatment of cognitive impairment in today’s society is increasing, and it is particularly imperative to take active intervention measures.

The process of aging is often accompanied by cognitive decline which eventually may lead to mild cognitive impairment (MCI) or AD[3]. The prevalence of MCI was shown to increase with age, and the incidence of further development of AD in MCI patients older than 65 years was 14.9%[4]. There are no drug can completely cure cognitive impairment in clinical practice,it is imperative to prevent senile cognitive impairment.To prevent the appearance of senile cognitive impairment is consistent with the viewpoint of “preventive treatment of disease” in traditional Chinese medicine guided by the holistic concept focuses on overall regulation.

Mitochondria, as the main "energy factory" of the body, provide most energy needed for life activities. In the process of gradual aging, mitochondrial function declines and the accumulation of dysfunctional mitochondria gradually affects the normal physiological functions of the body. Mitophagy is a self-regulatory phenomenon in response to adverse external stimuli, which can selectively remove abnormal cells and organelles in the body, and is critical for the normal physiological function[5]. The regulation of mitochondrial autophagy can eliminate dysfunctional mitochondria in cells, which can also promote cell differentiation, delay aging, and improve cognitive impairment[6]. The PINK1/Parkin mediated signaling pathway is one of the critical pathways in mitophagy. When mitochondria are damaged, Pink1 accumulates in the outer membrane of mitochondria due to transfer blockage, and Parkin is phosphorylated and activated after binding with the accumulated Pink1, thus initiating the signaling pathway. In recent years, more and more studies have shown that activating mitophagy can improve cognitive impairment caused by many reasons[7–8]. In this study, we investigated the effects of "Shuanggu Yitong" electroacupuncture (EA) pretreatment on cognitive impairment, mitochondrial function, and mitophagy in aging rats, and analyzed the potential mechanisms.

## Materials and Methods

### Animal grouping and model construction

Forty male SD rats (3 months old) were randomly divided into blank group, model group, EA group, and sham EA group (n=10 rats in each group). After adaptive feeding for 3 days, except for SD rats in the blank group, the rats in model group, EA group, and sham EA group were intraperitoneally injected with D-galactose solution with a concentration of 20 mg/mL for 40 days, according to the administration standard of 125 mg/kg/d, for the construction of subacute aging models. In the blank group, the same volume of 0.9% normal saline was injected according to the body weight.

The blank group and the model group: the rats were taken in the same time points as the EA group and the sham EA group, and then hung on an iron rack for fixation after rat clothes were worn to prevent them from moving. However, acupuncture and electroacupuncture were not performed[9].

EA group: at the beginning of model construction, the three acupoints of Baihui (DU20), Zusanli (ST36) and Guanyuan (CV4) were selected according to the textbook "Subject of experimental acupuncture and moxibustion", and the acupoint areas were shaved in advance for treatment. The rats wore self-made clothes to prevent movement, and were hung on an iron rack. After the skin surface of the acupoints was routinely disinfected with 75% alcohol, the rats were generally acupunctured with a filiform needle (Hwato brand, specification 0.30 mm × 13 mm). The specific needling depth was 2-3 mm for Guanyuan (CV4) and both sides Zusanli (ST36), and the two points were directly needled. Subsequently, the HANS LH202H-type electroacupuncture instrument was connected. Continuous wave mode was set, with a frequency of 2 Hz and a current of 1 mA, and it was powered on for 20 minutes. The electroacupuncture instrument was alternately connected to Guanyuan (CV4) and Zusanli (ST36) on both sides. The Baihui (DU20) was needled horizontally at a depth of 2 to 3 mm. To prevent rats biting, the electroacupuncture instrument was not connected at this acupoint. The operation was performed once every other day for a total of 8 weeks.

Sham EA group: At the beginning of model construction, the three acupoints of Baihui (DU20), Zusanli (ST36) and Guanyuan (CV4) were selected according to the textbook "Subject of experimental acupuncture and moxibustion", and the acupoint areas were shaved in advance for acupuncture. The rats wore self-made clothes to prevent movement, and were hung on an iron rack. After, the skin surface of the acupoints was routinely disinfected with 75% alcohol. Then, the filiform needles (Huaato brand, specification 0.30 mm × 13 mm) were fixed under the skin of the acupoints but not penetrating, and the electroacupuncture instrument was connected without current. The operation was also performed once every other day for a total of 8 weeks.

### Morris Water Maze Test

Morris water maze test was used to detect the cognitive function of the rats. The classic Morris water maze test procedures mainly included positioning navigation test and spatial exploration test. The detailed procedures were the same as described in the literature.

### HE Staining

The morphological changes of hippocampus in different groups of rats were observed by HE staining. The paraffin-embedded slices were baked in an oven at 60°C for 1-2 hours, and dewaxed with xylene for 5 min. They were then dyed with hematoxylin for 5-7min, and rinsed with tap water to remove residual color. Subsequently, they were treated by 0.1% ethanol hydrochloride for 2-5s, and immersed with tap water for 30s. Next, they were dehydrated using 100% ethanol for 30 seconds. Subsequently, they were treated by xylene and sealed with neutral gel. The hippocampal tissues were then observed under a microscope.

### Nissl Staining

Nissl staining was used to observe changes in the number of hippocampal neurons in rats. The paraffin-embedded slices were dewaxed by xylene for 15 min, and washed by distilled water for 30 seconds. The slices were stained by Toluidine blue dye solution at 60℃ for 45 min, and then washed by distilled water for three times. After that, they were treated by 95% ethanol until Nissl bodies were clearly observed under a microscope. Then they were dehydrated by 100% ethanol rapidly, and treated by xylene. Finally, sealing was made with neutral gel.

### Flow Cytometric Analysis

Mitochondrial membrane potential assay kit (C2006, Beyotime Biotechnology) was used to detect mitochondrial membrane potential (MMP). The main experimental instrument was a LX analytical flow cytometer (CytoFLEX LX), the instrument brand was Beckman, and the detection procedures were performed based on the kit instructions.

Mitochondrial membrane permeability transition pore (MPTP) was detected by flow cytometry, and the Mitochondrial Permeability Transition Pore Assay Kit (C2009S, Beyotime) was used. Sample processing and detection were carried out according to the kit instructions.

### Activity Detection of Respiratory Chain Complex I

The BC0510 Mitochondrial Respiratory Chain Complex I Activity Assay Kit produced by Solarbio was used to detect the activity of respiratory chain complex I. The detection procedures were carried out according to the kit instructions.

### Western blotting (WB)

The expression of PINK1 and Parkin in the hippocampus of the rats in different groups was determined by WB. The reagents used included: RIPA lysate (P0013B, Beyotime), PMSF (100 mM, BL507A, biosharp), phosphorylated protease inhibitor (P1081, Beyotime), Peroxidase AffiniPure Goat Anti-Rat IgG (H+L) (112-035-003, Jackson), Peroxidase AffiniPure Rabbit Anti-Goat IgG (H+L) (305-035-003, Jackson), ECL chemiluminescent substrate (BL520A, biosharp).

### Data Processing and Statistical Analysis

Continuous data were expressed as mean ± standard deviation, and the SPSS 19.0 software was used for statistical analysis. Independent-sample

*t*-test was used to compare the mean between two groups. *P* < 0.05 indicated significant difference.

## Results

### “Shuanggu Yitong” EA pretreatment improved the cognitive impairment in subacute aging rats

The results of the positioning navigation experiment are shown in Figure 1. The rats in the blank group showed a clear positioning of the platform in the fourth quadrant, and swam around the platform until they climbed it up. The rats in the model group mainly moved around the inner wall of the pool, and finally did not find the location of the platform. In the EA group, the rats mainly moved blindly in the center of each quadrant, while the sham EA group moved even more blindly than the EA group.

**Figure 1.**
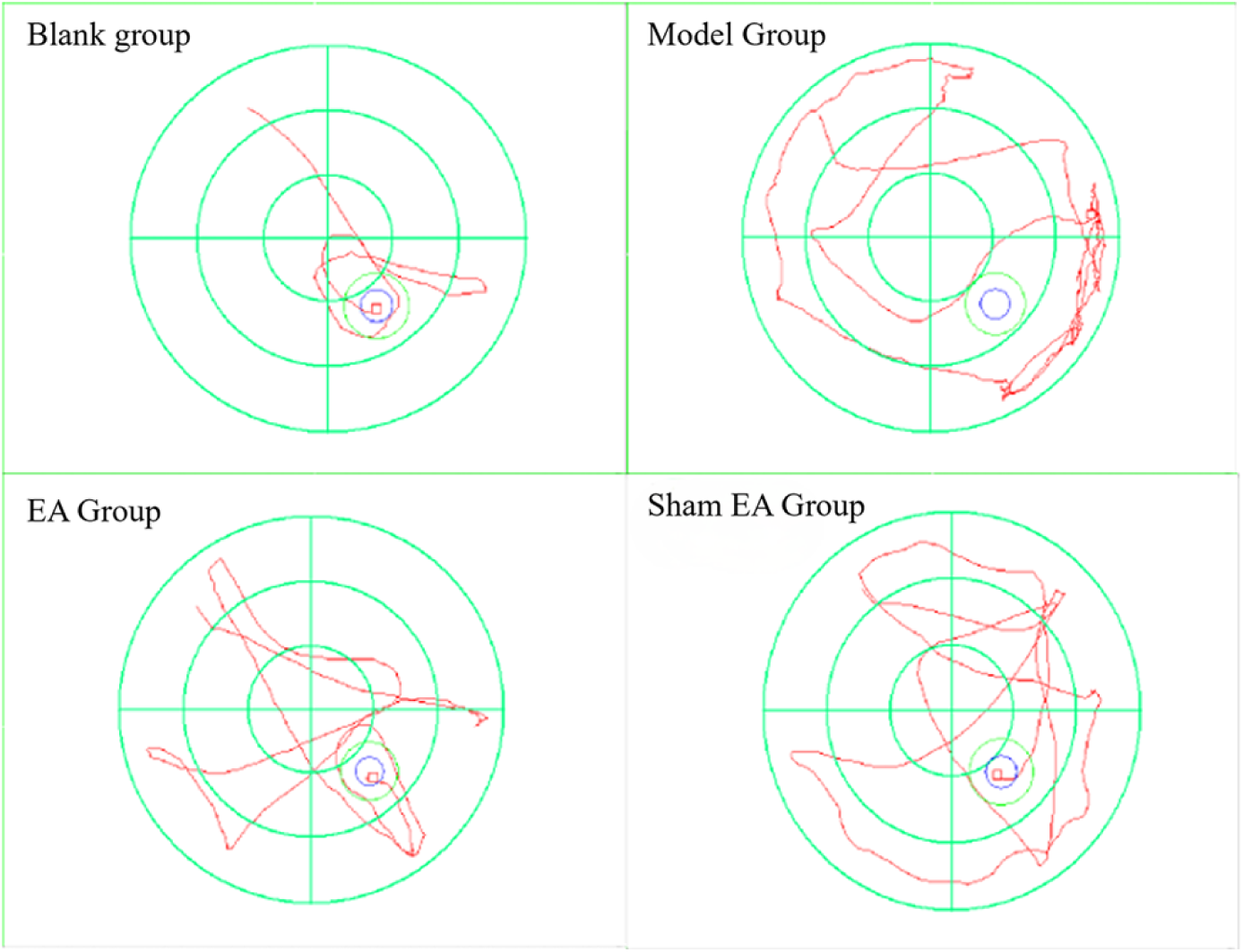
Trajectory diagram of Positioning Navigation

It can be seen from Table 1 that, compared with the blank group, the escape latency of the rats in the model group was significantly increased (*P* < 0.01) during the 5 days of the positioning navigation test, indicating that the cognitive impairment of the model group was more obvious, and that the model of subacute aging rats was successfully constructed. Compared with the model group, the daily escape latency of the rats in the EA group was reduced (*P* < 0.05), indicating that the "Shuanggu Yitong" EA pretreatment improved the cognitive impairment of the subacute aging rats. Compared with the model group, the cognitive impairment of the sham EA group was not significantly changed.

**Table 1.** Escape latency of positioning navigation test.

| Group | Escape Latency (s) |  |  |  |  |
| --- | --- | --- | --- | --- | --- |
|  | Day 1 | Day 2 | Day 3 | Day 4 | Day 5 |
| Blank group | 54.28 ± 8.57 | 40.13 ± 7.26 | 25.15 ± 6.47 | 15.89 ± 5.31 | 11.26 ± 2.86 |
| Model Group | 71.08 ± 7.57** | 57.60 ± 9.96** | 44.21 ± 4.23** | 30.61 ± 9.61** | 23.11 ± 8.15** |
| EA Group | 59.64 ± 6.72* | 43.44 ± 7.24* | 34.67 ± 6.53* | 18.67 ± 4.79* | 13.76 ± 3.08* |
| Sham EA Group | 64.99 ± 7.91 | 51.25 ± 6.35 | 41.13 ± 8.79 | 25.15 ± 7.93 | 17.98 ± 5.41 |

\* $P < 0.05$ , compared with the model group; \*\* $P < 0.01$ , compared with the blank group.

The results of the spatial exploration test are shown in Figure 2. The activity trajectories of the rats in the blank group were concentrated in the fourth quadrant (target quadrant), and the number of times they crossed the platform was high. The rats in the model group mainly moved around the inner wall of the pool and blindly explored it, and the number of times they crossed the platform was very small. The rats in the EA group tended to move in the target quadrant and near the platform, and crossed the platform for many times. Compared with the EA group, the sham EA group showed more blind activities in a wider range, and crossed the platform for less times.

**Figure 2.**
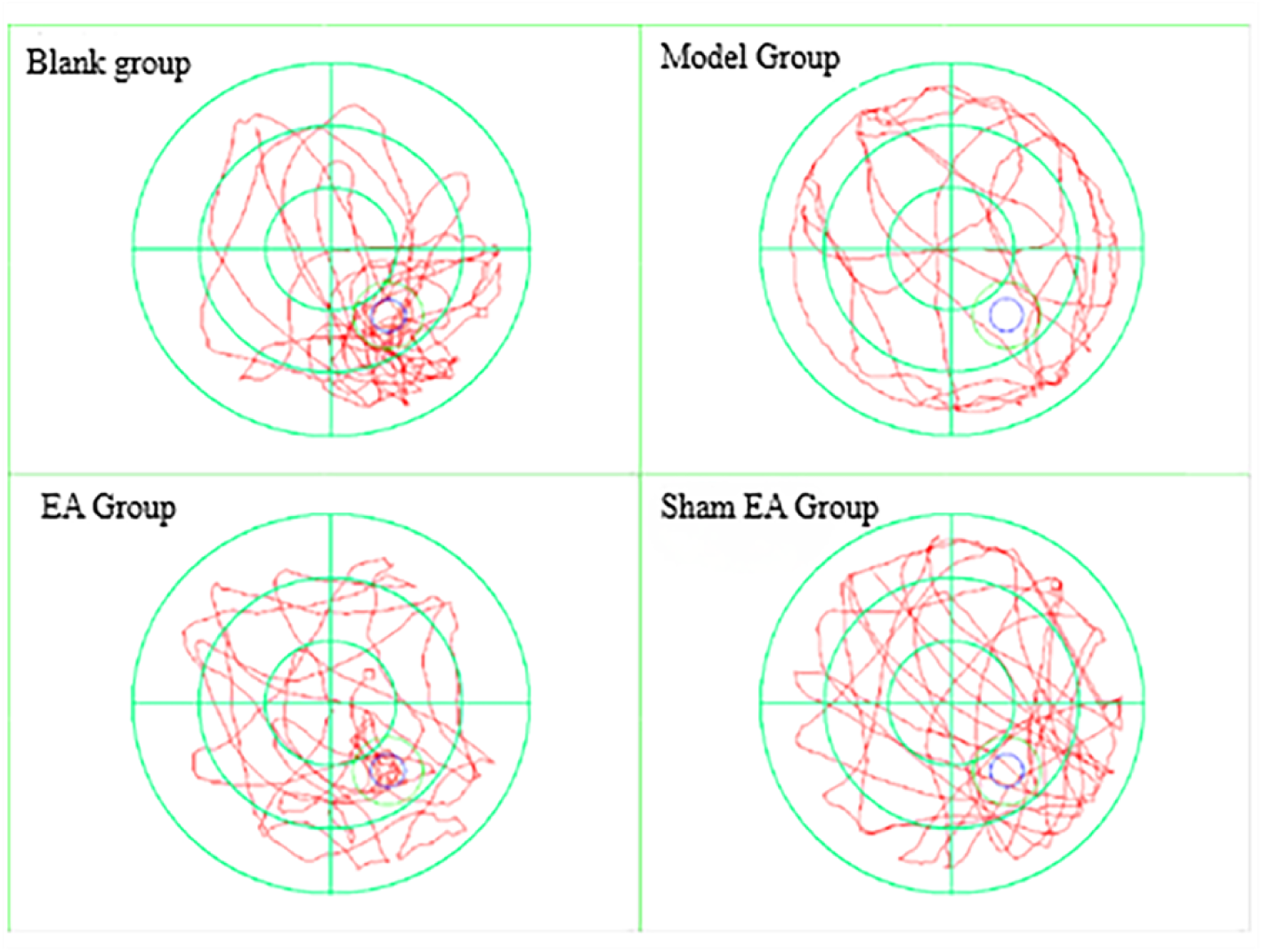
Trajectory diagram of spatial exploration test

It can be seen from Figure 3 that, compared with the blank group, the times of crossing the platform and the time of staying in the target quadrant of the rats in the model group were significantly reduced (*P* < 0.05), indicating that the cognitive impairment of the model group was more obvious, and that the model of the subacute aging rats was successfully constructed. Compared with the model group, the number of times of crossing the platform and the time of staying in the target quadrant in the EA group increased (*P* < 0.05), indicating that "Shuanggu Yitong" EA pretreatment improved the cognitive impairment of subacute aging rats. Compared with the model group, the sham EA group showed no significant improvement in cognitive impairment.

**Figure 3.**
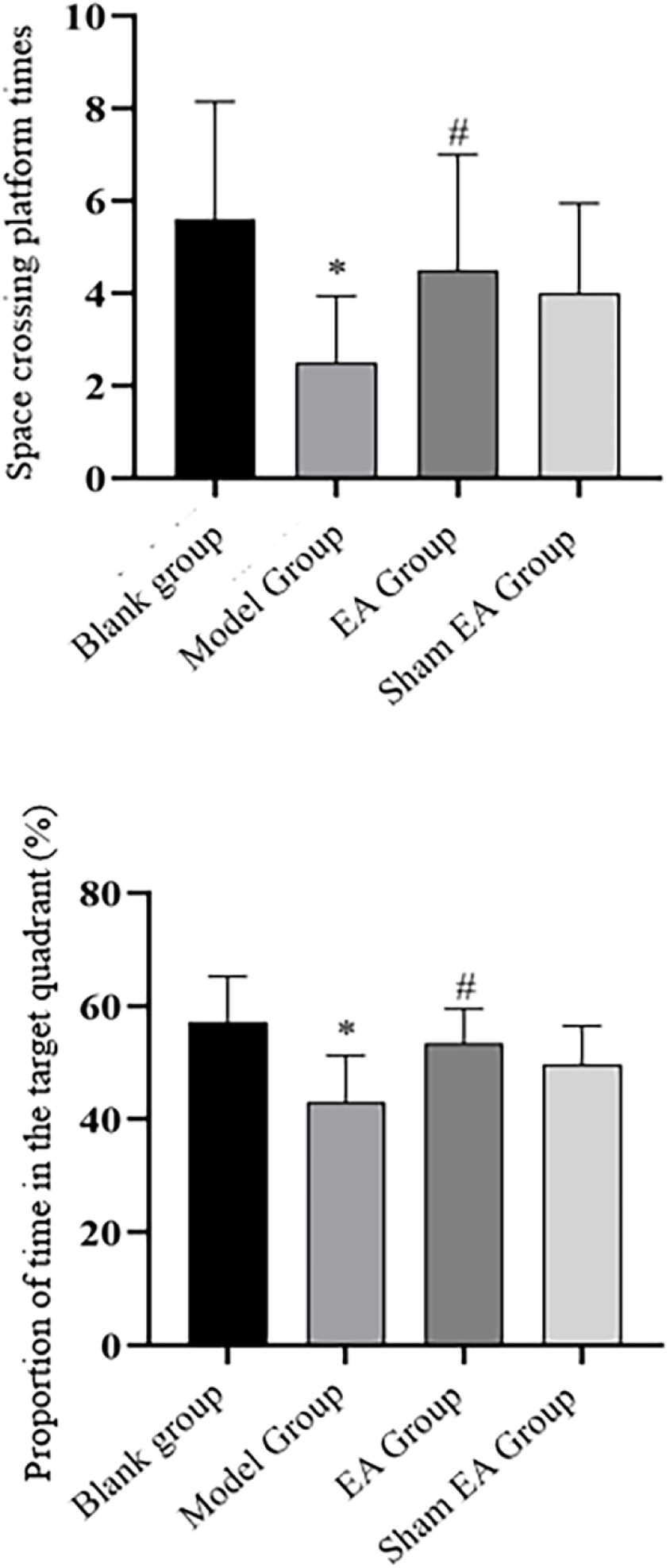
Space crossing platform times and proportion of time in the target quadrant of rats in each group. * *P* < 0.05, compared with the blank group; # *P* < 0.05, compared with the model group.

The same part of hippocampal CA1 tissues of rats in each group was selected for microscopic examination, and the results are shown in Figure 4. In the blank group, the neurons were arranged regularly and neatly, and the neuronal granules were plump, without injury and reduction. In the model group, the neurons were arranged disorderly with different degrees of injury and reduction (indicated by the black arrow), and some neurons were swollen and karyopyknosis was observed (blue arrow). Some neuron nuclei were even cracked into fragments (green arrow). Compared with the model group, the reduction of neurons in the sham EA group was improved (black arrow), and the neurons were recovered, but there were still some karyopyknosis (blue arrow) and karyorrhexis (green arrow), and the arrangement was still somewhat disordered. Compared with the model group, the neurons in the EA group were significantly improved, and the morphology of the neurons was similar to that of the blank group, with full granules and orderly arrangement, but there were still mild karyorrhexis (green arrow) and karyopyknosis (blue arrow).

**Figure 4.**
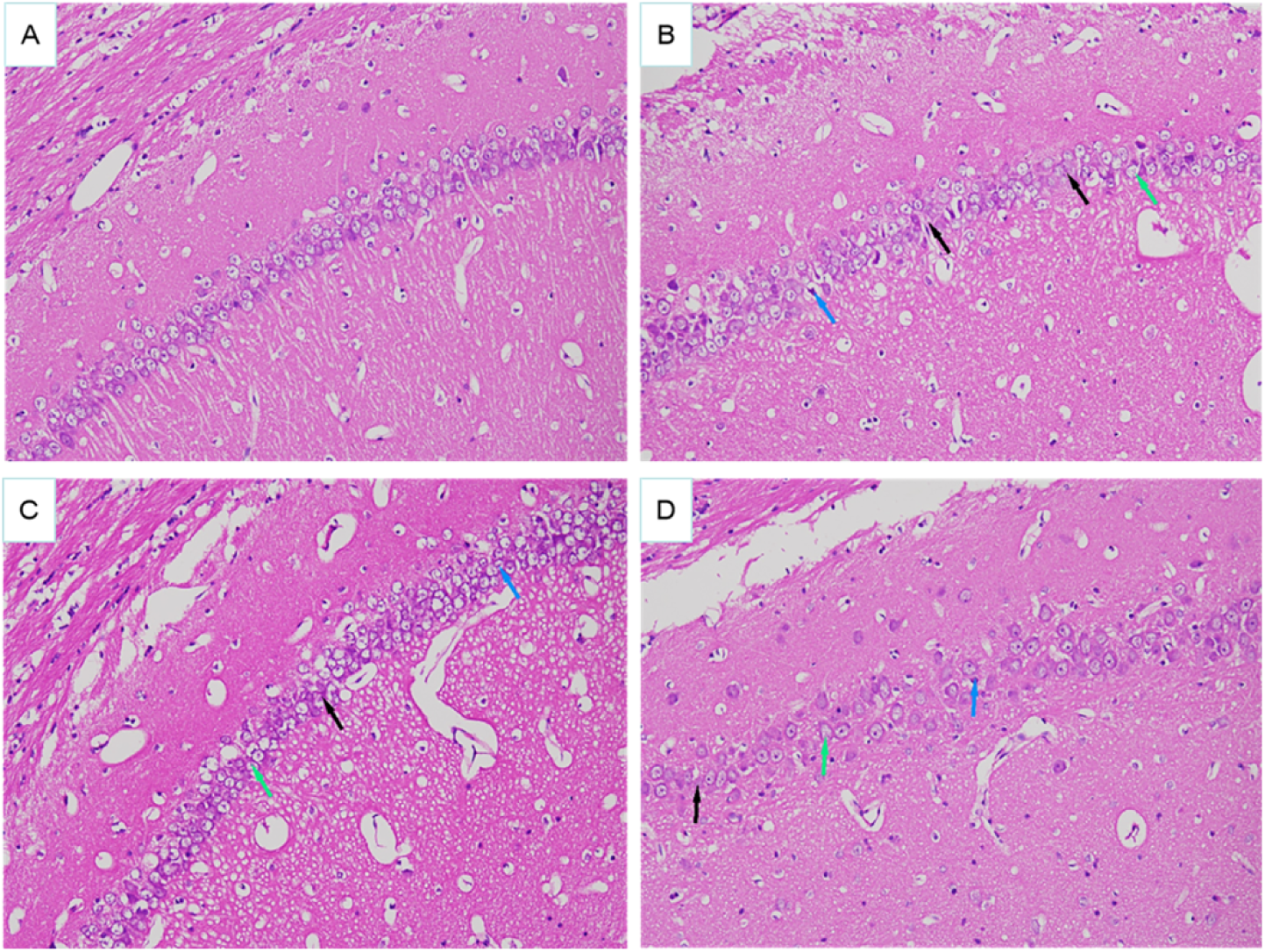
Morphological changes of hippocampal tissues in model rats (HE staining). HE staining results (×200). A: blank group; B: model group; C: sham EA group; D: EA group.

The same part of hippocampal CA1 tissues of the rats in each group was selected for microscopic examination, and results are shown in Figure 5. Compared with the blank group, the number of Nissl bodies and nuclei in the hippocampal CA1 region of the rats in the model group was significantly reduced, with irregular arrangement, and the staining was mild and vague. Compared with the model group, the number of Nissl bodies and nuclei in the hippocampal CA1 region of the rats in the sham EA group and the EA group was increased, with relatively regular arrangement, and the staining was deep and clear.

**Figure 5.**
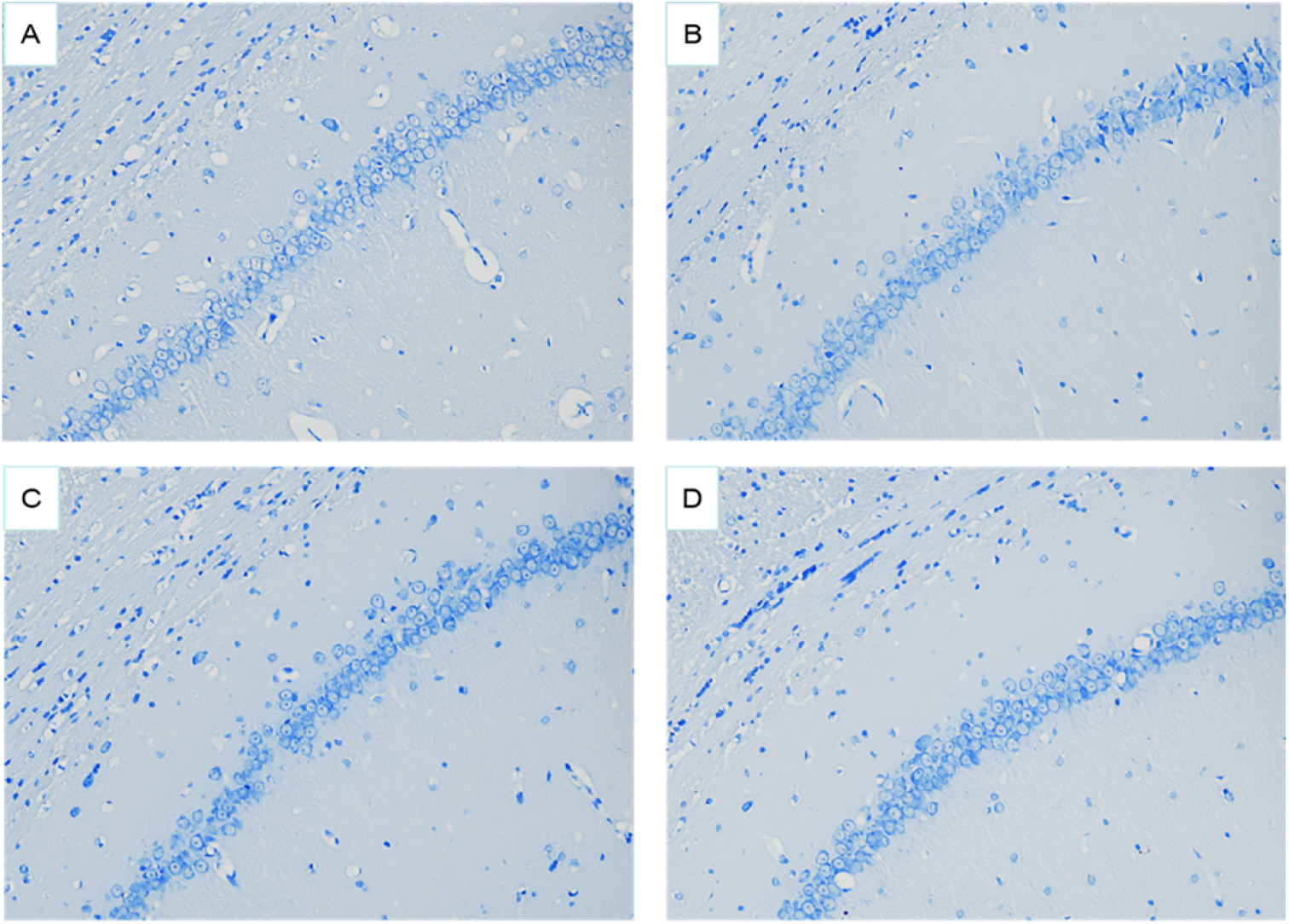
Changes in the number of hippocampal neurons in rats (Nissl Staining). Nissl staining results (×200). A: blank group; B: model group; C: sham EA group; D: EA group.

It can be seen from Table 2 that, compared with the blank group, the number of neurons in the hippocampal CA1 region of the rats in the model group was decreased significantly (*P* < 0.05), indicating that the number of neurons decreased gradually during the aging process. Compared with the model group, the number of neurons in the hippocampal CA1 area of the rats in the EA group and the sham EA group was increased significantly (*P* < 0.05), indicating that after intervention, the number of neurons increased, and that the learning, memory, cognitive and other functions of the rats were improved. ^a^*P* < 0.05, compared with the blank group; ^b^*P* < 0.05, ^c^*P* < 0.05, compared with the model group It can be seen from Figure 6 that, compared with the blank group, the number of neurons in the hippocampal tissue of the rats in the model group was significantly increased (*P* < 0.05), indicating that the number of neurons was decreased continuously during the aging process with cognitive impairment. After sham EA and EA treatment, the number of neurons in the two groups was increased significantly compared with the model group (*P* < 0.05), and the number of neurons in the EA group was increased even more obviously, which indicated that "Shuanggu Yitong" EA pretreatment could reduce the apoptosis of hippocampal neurons in the aging process with cognitive impairment, improve cognitive impairment, and delay aging.

**Figure 6.**
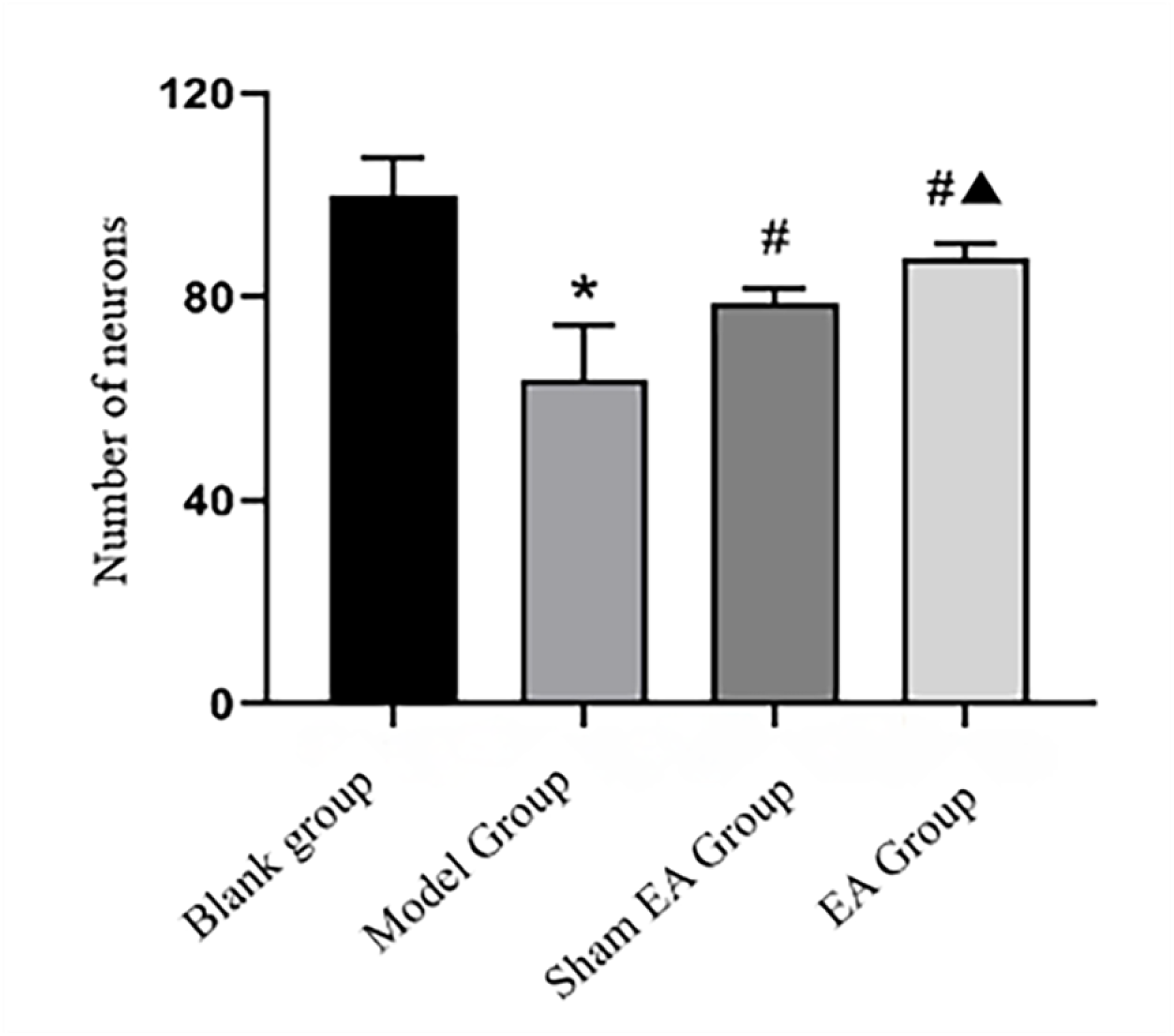
Comparison of the number of neurons in the hippocampus of rats in each group (X̄ ± S, 10 rats/group). * *P* < 0.05, compared with the blank group; #▲ *P < 0.05*, compared with the model group

**Table 2.**
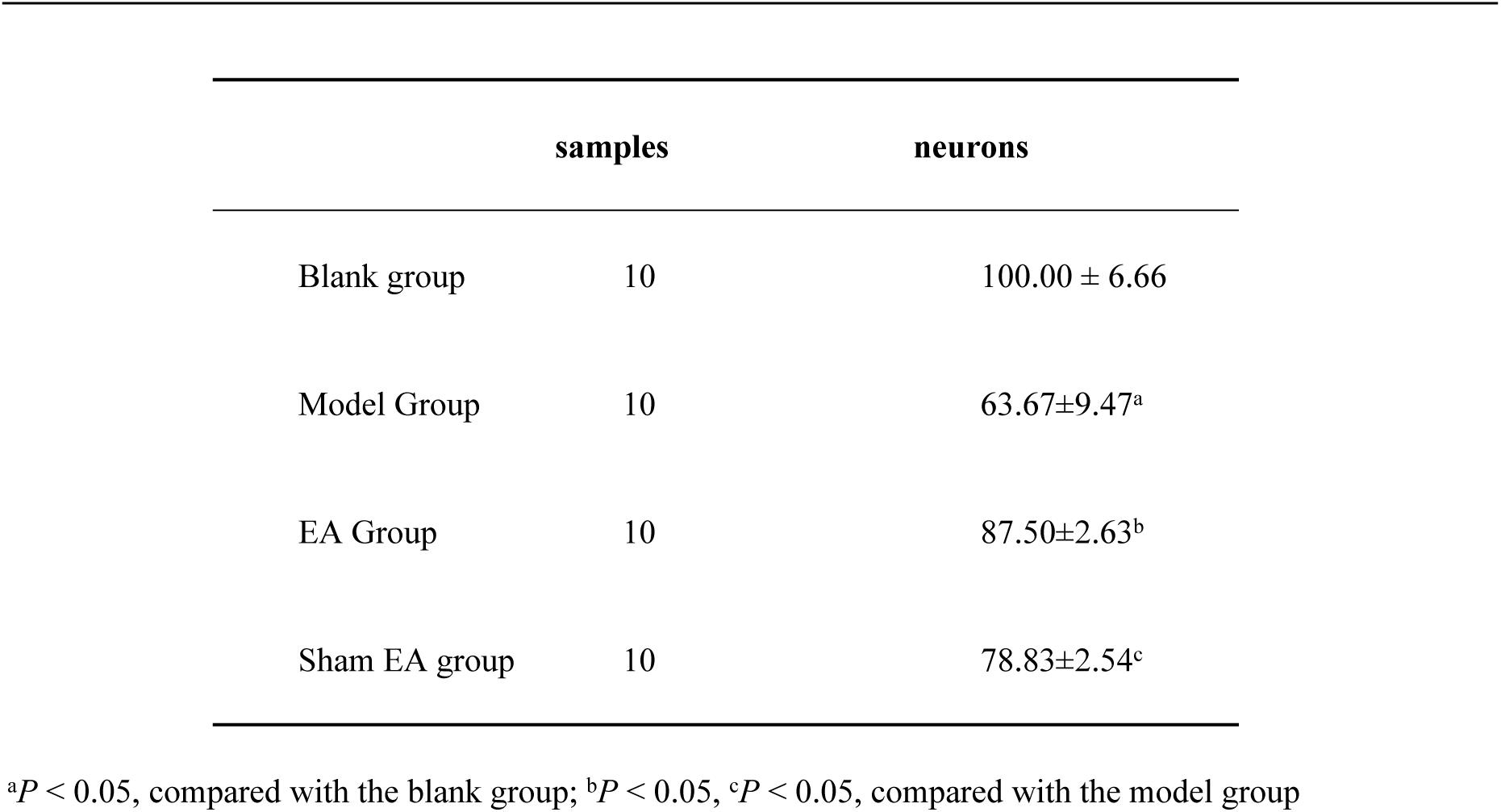
Number of Neurons (Microscope)

### Effects of "Shuanggu Yitong" EA Pretreatment on Mitochondrial Function and Mitophagy

Flow cytometry was used to measure MMP. As shown in Figure 7, the results showed that, compared with the blank group, MMP of rats in the model group was significantly depolarized (*P* < 0.01), indicating that MMP disappeared rapidly during aging. Compared with the model group, the depolarization of MMP in the EA group recovered after EA treatment (P < 0.05), indicating that the EA intervention during the aging process slowed down the disappearance of MMP and improved mitochondrial impairment.

**Figure 7.**
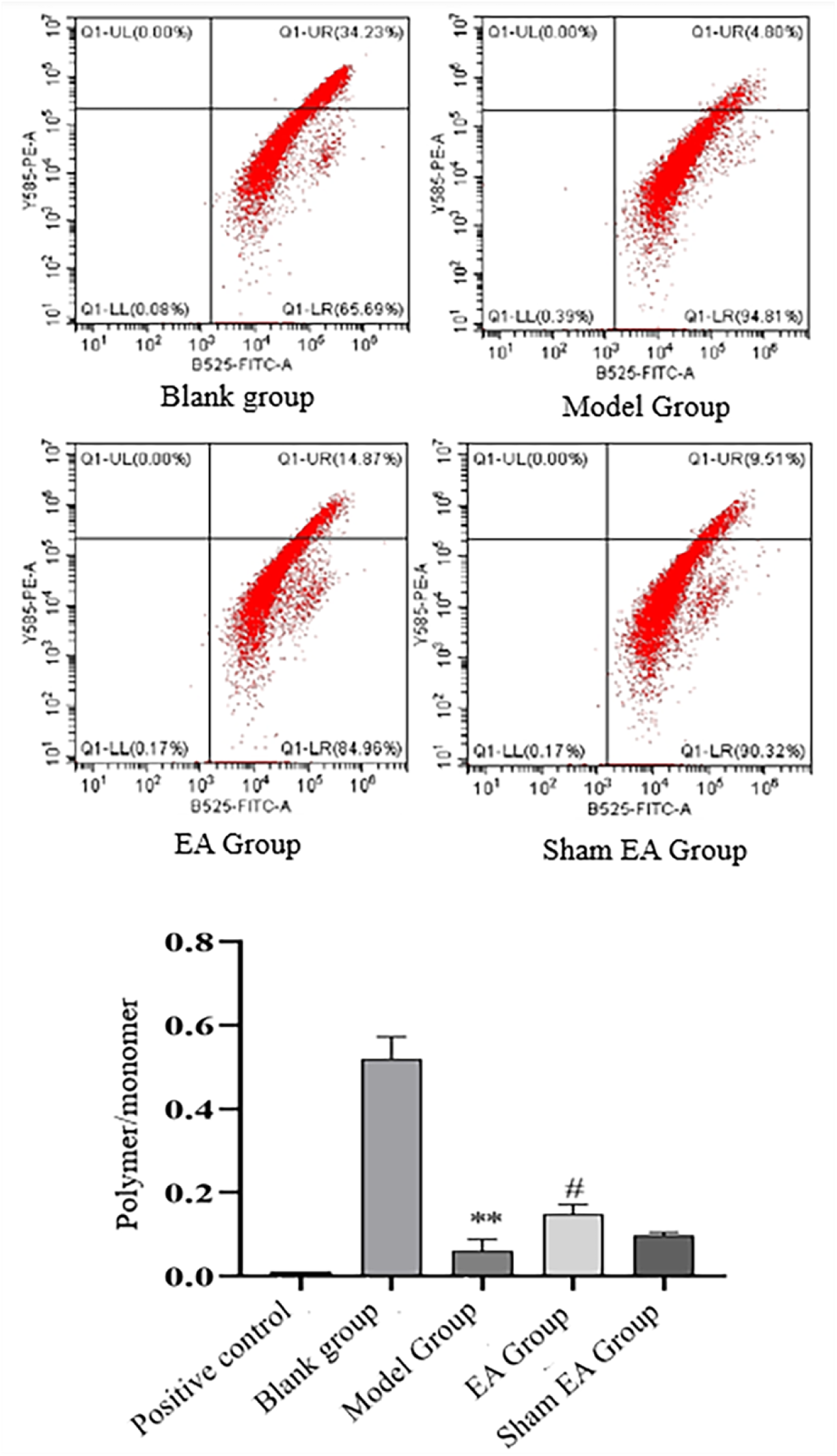
Measurement results of MMP in rats of each group (X̄ ± S, 10 rats/group). ** *P* < 0.01, compared with the blank group; # *P* < 0.05, compared with the model group.

As shown in Figure 8, the respiratory chain complex I activity test results revealed that complex I was mainly observed in the supernatant, and rarely found in the precipitate, which had mild effect on the experimental results. Compared with the blank group, the activity of complex I in the supernatant of the model group was significantly decreased (*P* < 0.01), and the total activity of complex I was also decreased (*P* < 0.05), indicating that the activity of complex Ⅰ was gradually decreased during aging, the level of mitochondrial oxidative phosphorylation was gradually decreased, and the normal function of mitochondria was reduced. Compared with the model group, the activity of complex I in the supernatant of the EA group was significantly increased (*P* < 0.01), and the total activity of complex I was also increased (*P* < 0.05), indicating that after the intervention of "Shuanggu Yitong" EA in the aging process, the activity of mitochondrial complex I was increased, the level of mitochondrial oxidative phosphorylation was increased, and the mitochondrial impairment was improved.

**Figure 8.**
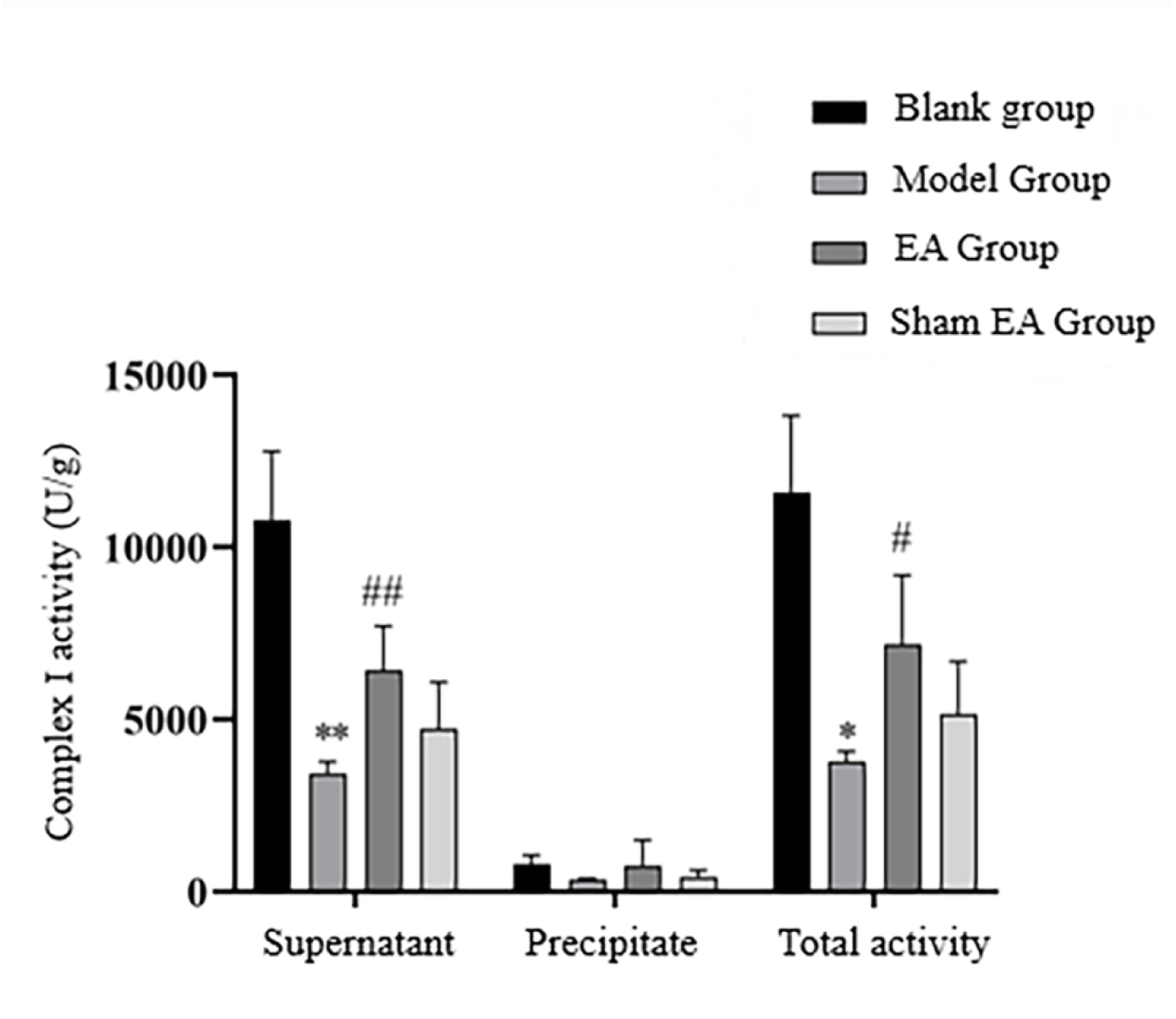
Measurement results of complex I activity in rats of each group (X̄ ± S, 10 rats/group). ** *P* < 0.01, ##*P* < 0.01, compared with the blank group; # *P* < 0.05, compared with the model group.

In addition, WB was used to detect the expression of PINK1 and Parkin in the hippocampus. The results are shown in Figure 9. Compared with the model group, the expressions of PINK1 and Parkin in the rats of the sham EA group were not significantly different (*P* > 0.05), indicating that the influence of physical stimulation interference factors from the acupuncture process on the rats was negligible. Compared with the blank group, the expressions of PINK1 and Parkin in the model group were significantly increased (*P* < 0.05), indicating that mitophagy was continuously enhanced during aging. After EA treatment, the expressions of PINK1 and Parkin in the EA group were significantly higher than those in the model group (*P* < 0.05), indicating that mitophagy was further activated after EA treatment, and mitochondria with abnormal function *in vivo* were cleared in time.

**Figure 9.**
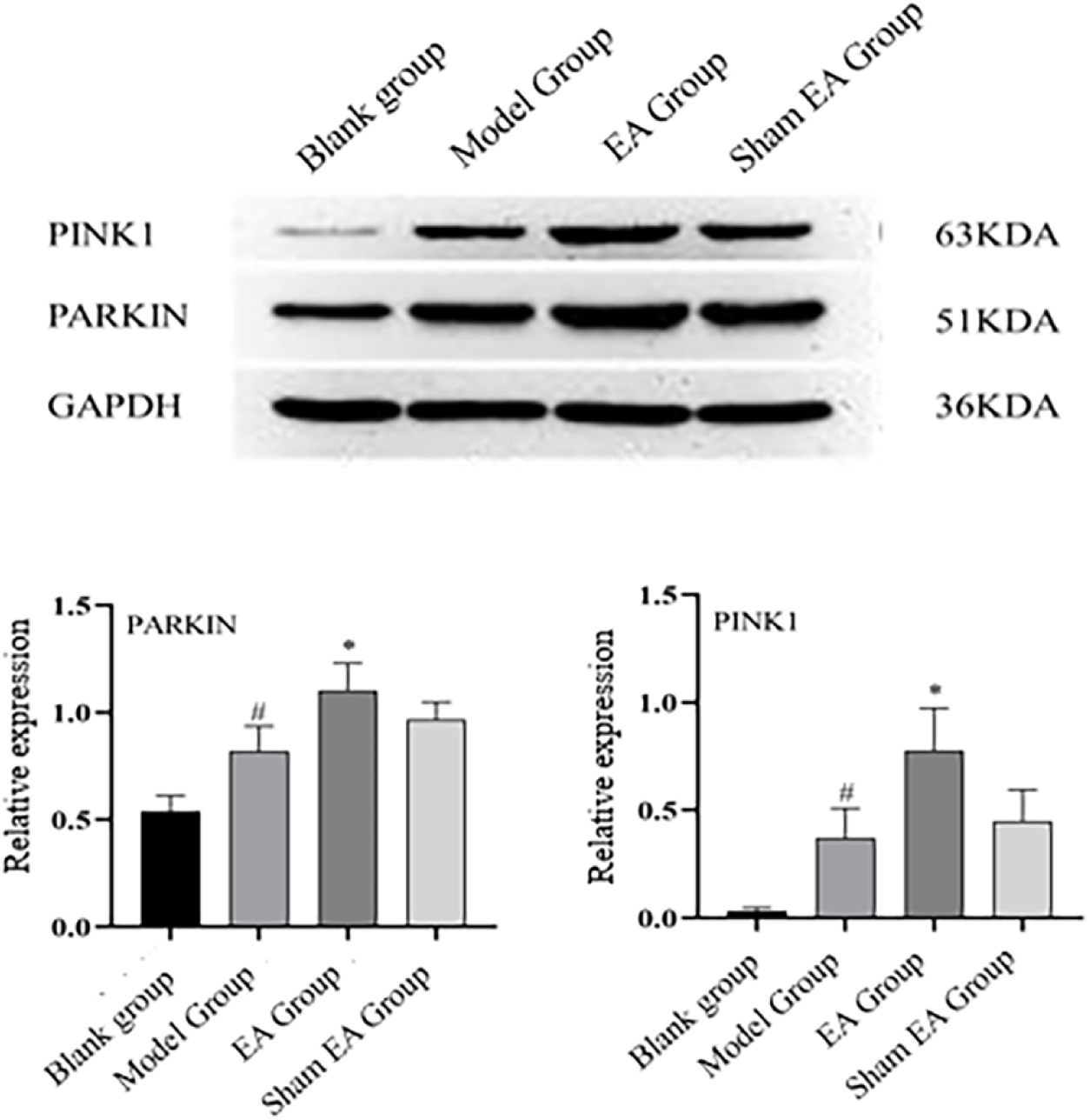
Comparison of PINK1 and Parkin protein expressions in the hippocampus of rats in each group (X̄ ± S, 10 rats/group). # *P* < 0.05, compared with the blank group; * *P* < 0.05, compared with the model group.

The results of MPTP test are shown in Figure 10. Compared with the blank group, MPTP opening of rats in the model group was increased significantly (*P* < 0.05), indicating that MPTP opening was increased gradually during aging. Compared with the model group, opening of MPTP in the EA group was significantly increased (*P* < 0.05), indicating that opening of MPTP was further increased after EA pretreatment.

**Figure 10.**
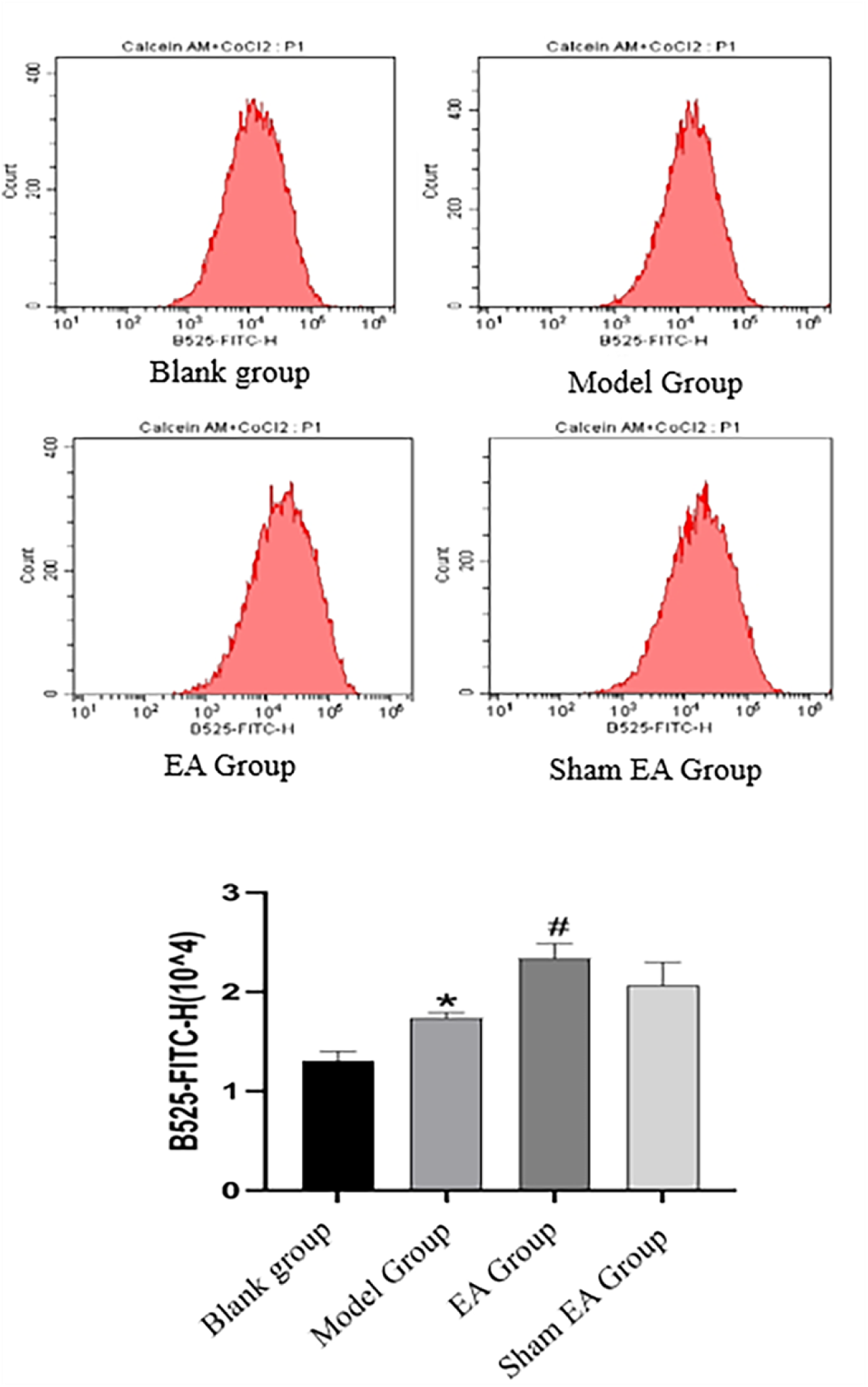
MPTP Flow Cytometer results of rats in each group.

## Discussion

Neurodegenerative diseases, including AD, Parkinson’s disease, amyotrophic lateral sclerosis, are unsolved disorders in the process of human aging. Clinically, there are no drugs available for the cure of neurodegenerative diseases, and some existing treatments can only alleviate or inhibit such disorders[10–11]. Neurodegenerative diseases have become a category of disorders that need to be solved urgently in the world. It is urgent for the medical scientists and human society to enhance research on the mechanism of neurodegenerative diseases and find more effective treatment modalities.

Hippocampus is critical for the learning, memory and cognition functions, which is closely related to the development of neurodegenerative diseases. Adult neurogenesis occurs in this region, which reflects plasticity of the brain in a sense[12]. Therefore, enhancement of adult hippocampal neurogenesis may promote treatment of neurodegenerative diseases[13]. Hippocampus, located at the base of the medial temporal lobe, is an important part of the limbic system, which is composed of several critical cortical areas, including parahippocampal gyrus, dentate gyrus, etc. Parahippocampal gyrus mainly includes CA1, CA2, CA3, and CA4. Parahippocampal gyrus is mainly composed of some pyramidal neurons, and critical for learning and memory. Parahippocampal gyrus is related to continuous self-renewal of neural stem cells, which affect hippocampus’s migration, proliferation, differentiation and maturation[14]. Nerves in hippocampus have received wide attention, because they can regenerate to replace old or damaged neurons with new neurons[15]. In this study, mitochondrial function and mitophagy of hippocampal CA1 neurons were detected to explore the possible mechanism of "Shuanggu Yitong" EA pretreatment improving senile cognitive impairment.

Autophagy is featured by intracellular degradation of cytoplasmic components in eukaryotes, which is a highly conserved degradation process of proteins or organelles. Autophagy is critical for maintaining metabolic balance of cells, and is a self-protection mechanism of cells[16]. In this process, some damaged proteins or organelles are packaged by autophagic vesicles with double-layered membrane structures. They eventually fuse with lysosomes and are degraded. The amino acids and other small molecules produced after degradation can be reused, which play an important role in maintaining cell homeostasis[17]. Autophagy and cell senescence are very important biological phenomena. Autophagy regulates the synthesis, degradation and reuse of intracellular substances, thus affecting all aspects of biological life processes[18].

In the process of aging, the functions of mitochondria and mitophagy are declining. The weakening of mitophagy can cause accumulation of damaged mitochondria in cells, reduce the efficiency of oxidative phosphorylation, increase reactive oxygen species in cells, and cause oxidative stress damage to cells, thereby further aggravating aging and forming a vicious circle[19]. A variety of substances are involved in the regulation of mitophagy in the process of delaying aging: the restoration of mitophagy through overexpression of parkin or inhibition of mammalian target of rapamycin can prevent mitochondrial dysfunction and aging[20]. Parkin can effectively resist stress-induced apoptosis and senescence of the bone marrow mesenchymal stem cells (BMSCs) by regulating mitophagy, and can improve the repair effect of BMSCs on early steroid-induced osteonecrosis of the femoral head[21]. PINK1 gene in rat bone marrow endothelial progenitor cells(BMEPCs) was knocked down by small interfering RNA technology, and it was found that down-regulation of PINK1 gene would aggravate aging of BMEPCs, suggesting that the aging and function of BMEPCs may be related to PINK1-mediated mitophagy[22]. Traditional Chinese medicine can delay aging by regulating mitophagy: acupoint catgut implantation can reduce the number of degenerated cells in the hippocampus and maintain the relatively normal structure of hippocampal tissues and neurons, which is related to the regulation of mitochondrial function and autophagy activity[23].

EA also participates in the regulation of autophagy. On the one hand, EA alleviates the corresponding diseases by inhibiting autophagy. EA can induce tolerance to cerebral ischemia by inhibiting autophagy through related signaling pathways, and can also inhibit hippocampal autophagy by reducing the expression of β-catenin/cyclooxygenase-2 protein, thereby inhibiting and alleviating central post stroke pain[24]. Sulforaphane can improve skeletal muscle fatigue induced by exhaustive exercise in mice by inhibiting mitochondrial autophagy mediated by PINK1/Parkin signaling pathway, which may be related to reducing skeletal muscle injury and improving antioxidant capacity and glycogen synthesis[25].On the other hand, EA can alleviate the corresponding diseases by promoting autophagy. EA may improve motor function in mice with spinal cord injury by promoting autophagy[26]. It also enhances autophagy in a polycystic ovary syndrome-like rat model to improve insulin resistance, mitochondrial dysfunction and endoplasmic reticulum stress[27]. Taken together, for acute diseases, although excessive autophagy under stress can protect the body from more damage, it causes unnecessary damage to the body, and EA can reduce the damage. For chronic diseases, the function of mitophagy is often weakened, resulting in accumulation of damaged cells in the body, which can not be removed in time. EA can promote removal of damaged cells and their organelles by autophagy in time, thus activating normal functions of the body.

In conclusion, "Shuanggu Yitong" EA pretreatment may timely remove abnormal mitochondria accumulated in the body during aging by enhancing PINK1/Parkin mediated mitophagy, thereby indirectly enhancing the overall normal function of mitochondria, so as to achieve the purpose of delaying aging and improving cognitive dysfunction.

## Author contributions

Biyong Liu and Tiantian Tan designed, performed and analysed the majority of experiments, and wrote the manuscript.

Su Qiu and Jialin Li, performed and analysed individual experiments.

Chengkai Xiong, Qing Liu and Yihong Li performed the data analysis and the formal analysis.

Jianmin Liu and Zhijie Li supervised the study, with contributions from all authors.

## Data Availability Statement

This is an open access article under the terms of the Creative Commons Attribution License, which permits use, distribution and reproduction in any medium,provided the original work is properly cited.

## References

[1]. Kanasi E, Ayilavarapu S, Jones J. The aging population: demographics and the biology of aging. Periodontol 2000. 2016;72(1): 13–18.

[2]. Grande G, Qiu C, Fratiglioni L. Prevention of dementia in an ageing world: Evidence and biological rationale. Ageing Res Rev. 2020; 64: 101045.

[3]. Morley JE. An Overview of Cognitive Impairment. Clin Geriatr Med. 2018;34(4):505–513.

[4]. Petersen RC, Lopez O, Armstrong MJ, et al. Practice guideline update summary: Mild cognitive impairment: Report of the Guideline Development, Dissemination, and Implementation Subcommittee of the American Academy of Neurology. Neurology. 2018; 90(3): 126–135.

[5]. Yang Y, Klionsky DJ. Autophagy and disease: unanswered questions. Cell Death Differ. 2020; 27(3): 858–871.

[6]. Yang JL, Mukda S, Chen SD. Diverse roles of mitochondria in ischemic stroke. Redox Biol. 2018; 16: 263–275.

[7]. Fang EF, Hou Y, Palikaras K, et al. Mitophagy inhibits amyloid-beta and tau pathology and reverses cognitive deficits in models of Alzheimer’s disease. Nat Neurosci. 2019; 22(3): 401–412.

[8]. Jiang W, Liu F, Li H, et al. TREM2 ameliorates anesthesia and surgery-induced cognitive impairment by regulating mitophagy and NLRP3 inflammasome in aged C57/BL6 mice. Neurotoxicology. 2022; 90: 216–27.

[9]. Liu Q, Xiong CK, Liu BY, et al. Improved Suspension Fixation Using Rat Jacket in Rat Acupuncture Experiments. J Vis Exp. 2023 Aug 18; (198). doi: 10.3791/65652.

[10]. Guo L, Fare CM, Shorter J. Therapeutic Dissolution of Aberrant Phases by Nuclear-Import Receptors. Trends Cell Biol. 2019; 29(4): 308–322.

[11]. Liu QQ, Ding SK, Shang YZ. Advance in the treatment of neurodegenerative diseases by traditional Chinese medicine. Journal of Chengde Medical University. 2021; 38(6): 518–521.

[12]. Wu S, Qing H, Liang JH. Advance in drugs that promote adult neurogenesis in the hippocampus. Acta Pharm Sin. 2016; 51(7):1025–1031.

[13]. Ding SK, Liu QQ, Liu XY, et al. Relationship between adult hippocampal neurogenesis and Alzheimer’s disease. Acta Neuropharmacologica. 2020; 10(6): 48–53.

[14]. Qiu CR Zhou BY, Wu ZL. Current research on the role of glutamate and its receptors in hippocampal learning and memory. Chin J Neuroanat. 2016; 32(4): 529–531

[15]. Wang JX, Hui Y, Gao X. Adult hippocampal neurogenesis and neurodegenerative diseases. Int J Genet. 2018; 41(3): 207–211.

[16]. Du CC, Liu LS, Li WJ, et al. Advances in autophagy regulating cell metabolic malance. Chinese Journal of Cell Biology. 2020, 42(11): 2003–2013.

[17]. Zhu Y, Liu X, Ding X, et al. Telomere and its role in the aging pathways: telomere shortening, cell senescence and mitochondria dysfunction. Biogerontology. 2019; 20(1): 1–16.

[18]. Birch J, Barnes PJ, Passos JF. Mitochondria, telomeres and cell senescence: Implications for lung ageing and disease. Pharmacol Ther. 2018; 183: 34–49.

[19]. Korolchuk VI, Miwa S, Carroll B, et al. Mitochondria in Cell Senescence: Is Mitophagy the Weakest Link? EBioMedicine. 2017; 21: 7–13.

[20]. Manzella N, Santin Y, Maggiorani D, et al. Monoamine oxidase-A is a novel driver of stress-induced premature senescence through inhibition of parkin-mediated mitophagy. Aging Cell. 2018; 17(5): e12811.

[21]. Zhang F, Wang L, Peng W, et al. p53 and parkin promote stem cells to repair osteonecrosis of the femoral head by regulating mitophagy to resist apoptosis and aging. Chin J Exp Surg. 2020; 37(11): 2015–2019.

[22]. Tang M, Shi Y, Xiao Z, et al. Effects of PINK1 mediated mitophagy on senescence and function of rat bone marrow endothelial progenitor cells. Chinese Pharmacological Bulletin. 2022; 28(10): 1472–1480.

[23 ]. Zhou M, Yuan Y, Lin Z, et al. Acupoint catgut embedding improves senescence in a rat model of ageing by regulating mitophagy via the PINK1 pathway. J Cell Mol Med. 2021; 25(8):3816–3828.

[24]. Zheng L, Li XY, Huang FZ, et al. Effect of electroacupuncture on relieving central post-stroke pain by inhibiting autophagy in the hippocampus. Brain Res. 2020; 1733: 146680.

[25]. Guo CD, Yang JX, Li PC. Reduction effect of sulforaphane on skeletal muscle injury and fatigue induced by exhaustive exercise through inhibiting mitochondrial autophagy mediated by PINK1/Parkin signal pathway. Chinese Journal of Food Hygiene. 2022; 34(6): 1158-1165.

[26]. Dai P, Huang SQ, Tang CL, et al. Effects of electroacupuncture at "Jiaji"(EX-B2) on autophagy and endoplasmic reticulum stress in spinal cord injury mice. Zhen Ci Yan Jiu. 2021; 46(1): 45–51.

[27]. Peng Y, Guo L, Gu A, et al. Electroacupuncture alleviates polycystic ovary syndrome-like symptoms through improving insulin resistance, mitochondrial dysfunction, and endoplasmic reticulum stress via enhancing autophagy in rats. Mol Med. 2020, 26(1): 73.

